# Temporal asymmetry of neural representations predicts memory decisions

**DOI:** 10.1101/2024.07.16.603778

**Authors:** Zhifang Ye, Yufei Zhao, Emily J. Allen, Thomas Naselaris, Kendrick Kay, J. Benjamin Hutchinson, Brice A. Kuhl

## Abstract

A stimulus can be familiar for multiple reasons. It might have been recently encountered, or is similar to recent experience, or is similar to ‘typical’ experience. Understanding how the brain translates these sources of similarity into memory decisions is a fundamental, but challenging goal. Here, using fMRI, we computed neural similarity between a current stimulus and events from different temporal windows in the past and future (from seconds to days). We show that trial-by-trial memory decisions (is this stimulus ‘old’?) were predicted by the *difference* in similarity to past vs. future events (temporal asymmetry). This relationship was (i) evident in lateral parietal and occipitotemporal cortices, (ii) strongest when considering events from the recent past (minutes ago), and (iii) most pronounced when veridical (true) memories were weak. These findings suggest a new perspective in which the brain supports memory decisions by comparing what actually occurred to what is likely to occur.

## Introduction

The ability to recognize a previously-encountered stimulus (recognition memory) is one of the most fundamental and well-studied forms of memory in both humans and non-human animals^1–3^. Over the past several decades, there has been substantial progress in identifying the brain regions that are involved in recognition memory decisions. In particular, univariate activation in subregions of lateral parietal cortex has been shown to scale with memory decisions (whether a stimulus is judged to be ‘old’ vs. ‘new’). However, a more elusive goal is to identify the specific computations that these brain regions perform in order to reach recognition memory decisions.

According to a highly influential class of computational models, recognition memory decisions are based on ‘global similarity’ (sometimes called ‘summed similarity’) between a current stimulus (a memory ‘probe’) and other recently-encountered stimuli. The core idea in these models is that if global similarity between the probe and recent experience is sufficiently high, the probe will be judged ‘old’^4–6^. These models, which are collectively referred to as global matching models, can explain an impressive number of findings from behavioral studies^7–9^. One particularly appealing aspect of these models is that they provide an elegant way of explaining why novel probes are sometimes falsely recognized. Namely, when a probe is novel, false recognition will occur if the probe has sufficiently high global similarity with other, studied stimuli.

To date, a few human fMRI studies have used pattern-based analyses to compute neural measures of global similarity. These studies have found that higher neural global similarity— including in lateral parietal cortex—is associated with a greater likelihood of endorsing a memory probe as ‘old’^10–12^. However, these studies suffer from a critical limitation: they do not consider the role of *time.* If neural measures of global similarity are capturing the influence that episodic memories of past experiences exert on current decisions, then time will be a critical factor. For example, events from the recent past should have a greater influence on current memory decisions than events from the distant past. However, it is alternatively possible that neural measures of global similarity do not, in fact, capture the influence of episodic memory but instead capture *time-invariant* effects of similarity. For example, a probe may have high neural similarity to other stimuli (whether they are in the past or even the future) simply because the probe is a more typical/common stimulus, or more consistent with schemas that have been generated from a lifetime of experience. This alternative account is important because it is well documented that when novel memory probes are more typical, they are more likely to be (falsely) judged as ‘old’^13,14^. Thus, to understand the neural computations that drive recognition memory decisions, it is imperative—but not trivial—to tease apart time-variant influences (e.g., recent experience) from time-invariant influences.

Here, in order to isolate the influence of recent experience on current memory decisions, we leveraged data from the Natural Scenes Dataset^15^—a massive human fMRI study in which 8 subjects each completed tens of thousands of trials of a continuous recognition memory test distributed over many months (Figure 1a, b). On each trial, subjects saw a natural scene image and decided whether the image was ‘old’ or ‘new’ (in the context of the experiment). On a trial-by-trial basis, we computed the fMRI pattern similarity of the current stimulus (probe) not only to events from the past (sampling from seconds to days in the past), but also to events in the future (the mirror image of events in the past). This unique analysis approach allowed us to identify brain regions that exhibited temporally-asymmetric relationships between global similarity and memory decisions. If memory decisions are more strongly influenced by neural similarity to past events compared to future events (i.e., a backward asymmetry), this provides unambiguous evidence for an influence of episodic memories on current decisions. Conversely, if memory decisions are driven by more generic effects of typicality (that are time-invariant), no temporal asymmetry would be expected.

**Figure 1.**
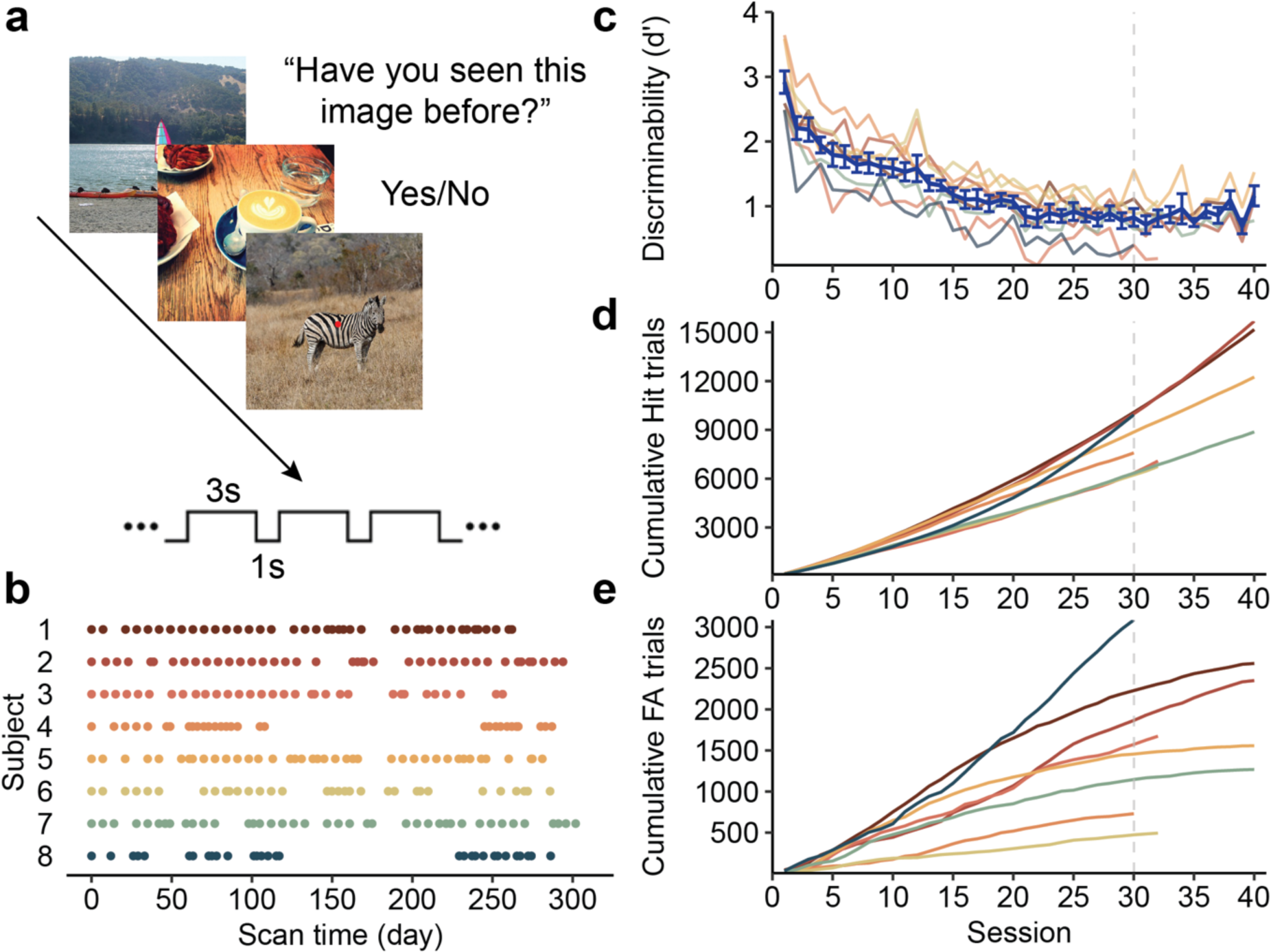
Experimental design and memory performance. **a,** Experimental design. Subjects performed a continuous recognition task on a series of natural scene images. On each trial, subjects indicated whether the current image had been presented at any point, so far, in the experiment. **b,** Task schedule. Each subject completed 30 – 40 fMRI scan sessions. The first session corresponds to day 0. **c,** Memory discriminability (d’) as a function of session number. Each colored line without error bars represents data from an individual subject. The blue line with error bars shows the mean d’ across subjects. Chance performance corresponds to a d’ of 0. The vertical grey dashed line marks the last session (30) included in the main analyses. **d**, The cumulative number of hit trials as a function of session number. **e,** The cumulative number of false alarm trials as a function of session number. Error bars reflect the standard error.

Motivated by numerous neuroimaging studies implicating lateral parietal cortex in recognition memory decisions^16–18^—and in representing the contents of memories^19–21^—we specifically predicted a backward asymmetry in lateral parietal cortex. That is, we predicted that the decision to endorse a probe as ‘old’ would be driven by the strength of lateral parietal similarity to past events *relative to* future events. For comparison, we also considered several additional regions of interest that are involved in memory, vision, and motor responses.

To preview, we show that recognition memory decisions are robustly predicted by backward asymmetry of global similarity in lateral parietal cortex. This influence was selective to events from the recent past (as opposed to more temporally-distant events) and was also related to the objective mnemonic history of a probe: global similarity had the strongest effect on memory decisions when the probe had not recently been encountered. Finally, using convolutional neural networks, we show that neural measures of global similarity that drive memory decisions primarily contain information about high-level semantic features. Collectively, these findings provide new insight into how recognition memory decisions are computed. In particular, our findings support an account of memory decisions in which time-variant similarity to recent events from the past is ‘baselined’ against time-invariant similarity (here, measured as similarity to future events).

## Results

### Recognition memory performance

Considering performance across all experimental sessions, mean recognition memory discriminability (d’) was 1.23 (range across subjects: 0.69 – 2.92), which was significantly above chance (*t*_(39)_ = 15.53, *p* < 0.001). However, performance significantly decreased over sessions (linear mixed-effects model, *χ*^2^_(1)_ = 308.04, *p* < 0.001) (Figure 1c). The mean hit rate across all sessions was 62.8% (54.6% – 86.5%) and the mean false alarm rate was 23.3% (4.6% – 39.9%). Linear mixed-effects models revealed that while the hit rate decreased across sessions (*χ*^2^_(1)_ = 74.35, *p* < 0.001), the false alarm rate increased (*χ*^2^_(1)_ = 117.76, *p* < 0.001).

One distinct advantage of the current data set is that it provides an incredibly large number of total trials per subject and, consequently, a very large number of both ‘hit’ trials (repeated images correctly identified as ‘old’) and ‘false alarm’ trials (novel images falsely identified as ‘old’). The mean number of hit trials per subject was 10,414 (range: 6749 – 15682) (Figure 1d) and the mean number of false alarm trials was 1,715 (range: 494 – 3,087) (Figure 1e).

Because not all subjects completed all 40 experimental sessions (range: 30 – 40 sessions), we restricted subsequent analyses to the first 30 sessions so that session effects were matched across subjects. Considering only the first 30 sessions, the mean d’, hit rate and false alarm rate were 1.34 (range: 0.78 – 2.92), 63.3% (range: 54.6% – 86.5%) and 20.2% (range: 4.6% – 32.5%), respectively. Across the first 30 sessions, each subject saw 9,209 novel images and 13,291 repeated images.

### Predicting memory decisions from neural global pattern similarity

Our overarching goal was to isolate the influence that past events exerted on memory decisions in the continuous recognition task. Because we hypothesized that the relative recency of past events would determine their influence, we separated past events into three temporal windows—immediate, recent, and distant—that corresponded to events from the same scan run (immediate), the same scan session (recent), or a different scan session (distant). Specifically, the *immediate temporal window* binned trials from the same scan run as the current probe, extending 15 trials in the past (mean temporal distance to probe = 35.0 seconds, range: 4.0 seconds to 68.0 seconds); the *recent temporal window* included trials from the preceding 3 scan runs, excluding trials within the same scan run as the probe (mean = 3.9 minutes, range: 2.8 minutes to 37.1 minutes); and the *distant temporal window* included trials from the prior fMRI session (mean = 7.3 days, range: 1.0 days to 28.0 days).

To measure the similarity of each memory probe to events from the past, we used fMRI pattern similarity to compute neural measures of *global similarity*. Specifically, for each memory probe, global similarity within a given brain region of interest (ROI) was obtained by taking the fMRI activity pattern from the current trial (probe) and correlating it (Pearson correlation) with the fMRI activity pattern for each of the trials within a given temporal window. These correlations were then Fisher z-transformed and averaged, yielding a global similarity value for each of the three temporal windows in the past. Critically, we also computed global similarity to stimuli *in the future* using the same approach and same three temporal windows, but for stimuli that had not yet been encountered. Finally, for each temporal window, we subtracted ‘forward’ global similarity (to future events) from ‘backward’ global similarity (to past events). All global similarity analyses reported below were based only on this difference score (Figure 2a). Our rationale for this approach was that any temporally-symmetric similarity effects would cancel out. For example, if a given scene image (probe) includes very common objects or landmarks, then it should be normatively similar to other scenes (whether they occurred in the past or the future). In contrast, any contribution of episodic memory to global similarity would necessarily be temporally asymmetric (past > future). Thus, subtracting forward similarity from past similarity is a simple, but powerful way to isolate the influence of past experience.

**Figure 2.**
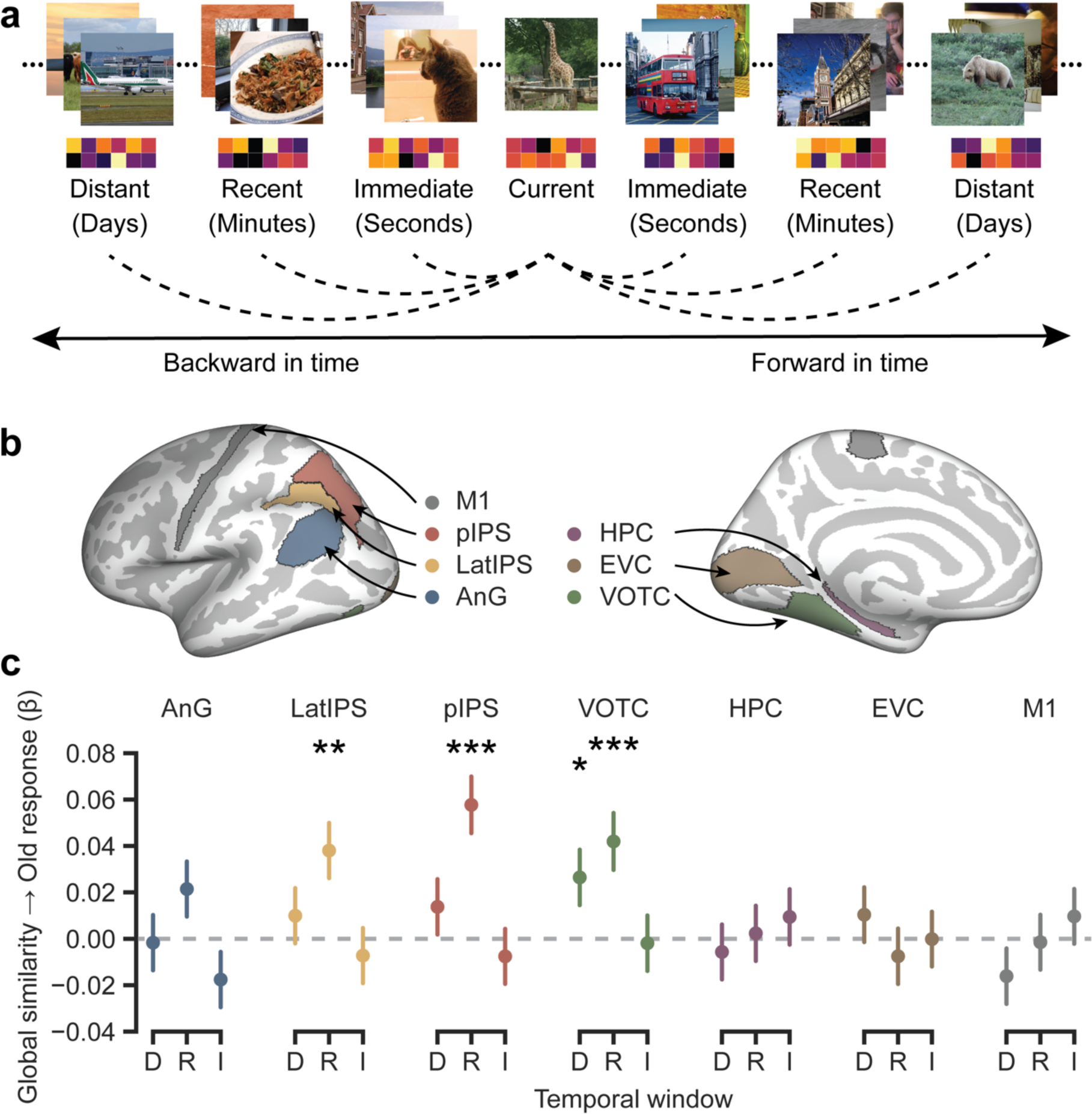
Global similarity effects. **a,** Global similarity of neural patterns was calculated between the current trial and trials within each temporal window. The global similarity values from the forward (in the future) temporal windows were subtracted from the corresponding global similarity values from the backward (in the past) temporal window. **b,** Regions of interest included angular gyrus (AnG), lateral intraparietal sulcus (LatIPS), posterior intraparieral sulcus (pIPS), ventral occipitotemporal cortex (VOTC), hippocampus (HPC), early visual cortex (EVC) and premotor cortex (M1). ROIs are illustrated on the inflated FreeSurfer fsaverage cortical surface. All ROIs were combined across the left and right hemispheres. **c,** The global similarity effects within each temporal window. A positive coefficient indicates that greater global similarity is associated with higher probability to endorse images as ‘old’. Error bar denotes standard error.

To test whether global similarity predicted memory decisions, we built mixed-effects logistic regression models in which global similarity values served as predictors and the outcome (dependent measure) was the memory decision for each probe (i.e., ‘old’ or ‘new’ response). Our initial models included global similarities from all three temporal windows as separate regressors. We also included a categorical regressor representing the veridical mnemonic history of the probe: whether the probe image was being presented for the 1^st^, 2^nd^ or 3^rd^ time (E1, E2, E3). In addition, session number and the proportion of old responses within each temporal window were also included in the model to account for potential decision criteria drift (across sessions) and the influence of response history (e.g., if a relatively high/low number of ‘old’ responses were made in a given temporal window). (See *Methods* for detailed model specifications).

Motivated by prior studies, we focused our fMRI analysis on lateral parietal cortex^10,11^, which we divided into three regions of interest (ROI): angular gyrus (AnG), lateral intraparietal sulcus (LatIPS), and posterior intraparieral sulcus (pIPS) (Figure 2b). We also included ventral occipitotemporal cortex (VOTC) given its role in representing the content of natural scenes images^22^ and the hippocampus (HPC) given its importance in episodic memory^23,24^. In addition, we included early visual cortex (EVC) and primary motor cortex (M1) as active control regions. For EVC, we reasoned that while it would represent low-level properties of currently-displayed stimuli, these representations would not be related to memory. For M1, we had no reason to expect it to be involved in representing scene content or to be related to memory, but it serves as a useful control given that it should track motor responses. Each ROI was associated with a unique mixed-effects logistic regression model.

As a first step, we tested for an omnibus global similarity effect by comparing full models (with all three global similarity regressors) to models without any global similarity regressors. This revealed that global similarity was predictive of memory decisions—with higher global similarity associated with a greater probability of an ‘old’ response—in LatIPS (*χ*^2^_(9)_ = 22.16, *p* = 0.008), pIPS (*χ*^2^_(9)_ = 31.75, *p* < 0.001), and VOTC (*χ*^2^_(9)_ = 27.61, *p* = 0.001), with all three models surviving correction for multiple comparisons. There was also a global similarity effect in EVC (*χ*^2^_(9)_ = 18.71, p = 0.028), that did not survive correction, and a trend toward an effect in AnG (*χ*^2^_(9)_ = 16.06, *p* = 0.066). There was no effect of global similarity in HPC (*χ*^2^_(9)_ = 2.83, *p* = 0.971) or M1 (*χ*^2^_(9)_ = 7.50, *p* = 0.585).

For the preceding analyses, all trials within a given temporal window were given equal weight (with pattern similarity simply averaged across all trials). While the idea of pooling across trials is central to global matching models, some models do give higher weight to past events that strongly match a current probe (i.e., high similarity matches)^25^. This does raise an important question of whether, in our analyses, there was any benefit to averaging across trials, as opposed to only using the most similar trials. Thus, we tested another set of models where, for each temporal window, we only included the similarity for the single trial that was most similar to the current probe. In other words, we replaced the averaged (global) similarity with the maximal similarity. For these models, regressors for each of the three temporal windows were included within the same model. Interestingly, maximal similarity did not predict memory decisions for any of the ROIs (AnG, *χ*^2^_(9)_ = 14.16, *p* = 0.117; LatIPS, *χ*^2^_(9)_ = 11.25, *p* = 0.259; pIPS, *χ*^2^_(9)_ = 13.69, *p* = 0.134; VOTC, *χ*^2^_(9)_ = 16.89, *p* = 0.051; HPC, *χ*^2^_(9)_ = 8.95, *p* = 0.442; EVC, *χ*^2^_(9)_ = 12.84, *p* = 0.170; M1, *χ*^2^_(9)_ = 6.24, *p* = 0.716). Thus, at least when comparing the extremes—averaging with equal weight (global similarity) vs. selecting the maximal similarity—there was a clear advantage to global similarity.

We next performed follow-up analyses again using global similarity to predict memory decisions, but separately for each temporal window (immediate, recent, distant). Interestingly, none of the ROIs exhibited a significant global similarity effect for the immediate temporal window (all *p’s* > 0.14), though it should be noted that this temporal window contained the fewest trials. For the recent temporal window, however, there were significant effects in LatIPS (*χ*^2^_(1)_ = 10.21, *p* = 0.001, survived correction), pIPS (*χ*^2^_(1)_ = 22.61, *p* < 0.001, survived correction), VOTC (*χ*^2^_(1)_ = 11.97, *p* < 0.001, survived correction) and a trend in AnG (*χ*^2^_(1)_ = 3.27, *p* = 0.071) (Figure 2c). There were no effects in EVC, HPC or M1 (*p’s* > 0.5). For the distant temporal window, only VOTC showed a significant global similarity effect (*χ*^2^_(1)_ = 4.90, *p* = 0.027), but it did not survive correction for multiple comparisons (all other regions: *p’s* > 0.18).

Taken together, the analyses thus far strongly implicate regions of lateral parietal cortex and VOTC in expressing representations that were predictive of memory decisions and specifically identify the recent temporal window—events that occurred minutes ago in the past—as being most influential. In subsequent analyses, we therefore focus on the lateral parietal and VOTC ROIs, and we restrict analyses to the recent temporal window.

### Influence of global similarity depends on mnemonic history of probe

In all of the global similarity models presented thus far, we included a regressor to account for the novelty of the probe—whether the probe was novel (1^st^ exposure; E1) or had been presented before (2^nd^ or 3^rd^ exposure; E2, E3). However, an interesting question is whether the influence of global similarity varies as a function of the novelty of the probe. In particular, we hypothesized that global similarity would have a relatively stronger influence on memory decisions for Novel probe trials compared to Old probe trials. Our rationale for this prediction was that when the probe was Novel, there is no ‘true’ memory signal (i.e., there is no event-specific true memory for the prior encounter) and, therefore, decisions would rely on global similarity (which pools across many trials). In contrast, for Old trials we reasoned that ‘true’ memory for a prior encounter with the probe would compete with—and largely override—the influence of global similarity. To test this, we constructed another set of mixed-effects logistic regression models. Here, based on results presented above, we only included the recent temporal window and only tested the lateral parietal ROIs and VOTC. Additionally, to create balance in the number of Novel vs. Old trials, we included E1 (1^st^ exposure; Novel) and E2 (2^nd^ exposure; Old) trials, but excluded E3 trials. Finally, and importantly, we excluded any E2 trials for which the corresponding E1 exposure fell within the recent temporal window. Thus, because E1 trials always occurred outside the temporal window from which global similarity was computed, the E1 trials did not directly contribute to global similarity values. As such, these analyses were not intended to test whether E1 trials had an effect on global similarity values; rather, the key question was whether E1 trials (that fell *outside* the global similarity window) weakened the *influence* of global similarity on memory decisions.

Across each of the lateral parietal and VOTC ROIs, there was a significant effect of global similarity on memory decisions for Novel trials (AnG, *β* = 0.089, *Z* = 4.34, *p* < 0.001; LatIPS, *β* = 0.072, *Z* = 3.52, *p* < 0.001; pIPS, *β* = 0.114, *Z* = 5.50, *p* < 0.001; VOTC, *β* = 0.074, *Z* = 3.52, *p* < 0.001; all survived correction) (Figure 3a). However, counter to our prediction, the effect of global similarity on memory decisions for Old trials was also significant in each of the lateral parietal and VOTC ROIs (AnG, *β* = 0.060, *Z* = 2.87, *p* = 0.004; LatIPS, *β* = 0.056, *Z* = 2.65, *p* = 0.008; pIPS, *β* = 0.080, *Z* = 3.79, *p* < 0.001; VOTC, *β* = 0.076, *Z* = 3.65, *p* < 0.001; all survived correction). Moreover, the global similarity effect was not significantly stronger for Novel trials than Old trials in any of the four ROIs (*p’s* > 0.24).

**Figure 3.**
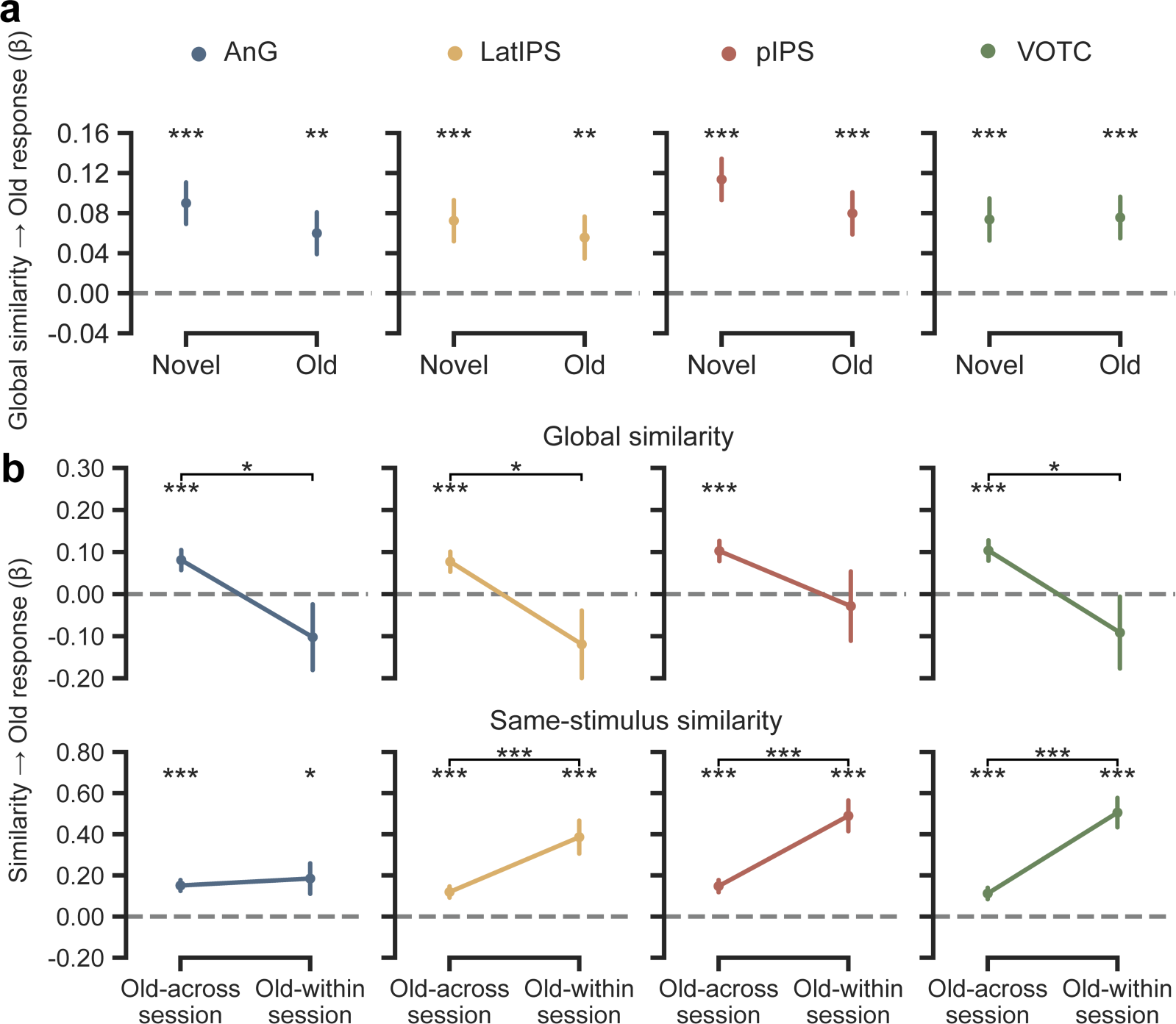
Global similarity effect as a function of mnemonic history. **a,** Global similarity effect on memory decisions for Novel (E1) and Old (E2) trials. **b,** Global similarity and same-stimulus effects for Old (E2) trials, separated as a function of when E1 occurred. Old-within trials are E2 trials for which the corresponding E1 trial occurred within the same experimental session. Old-across trials are E2 trials for which the corresponding E1 trial occurred in a prior experimental session. Error bar denotes standard error.

Although we predicted that global similarity would have a weaker effect on memory decisions when a ‘true’ memory signal was present (Old trials), one potential explanation why we did not see this effect is that, for many of the Old trials, a true memory signal may have been quite weak. Specifically, given the highly protracted nature of the experiment (analyses included 30 fMRI sessions per subject distributed over many months), for many of the Old trials (E2), the prior exposure of the stimulus (E1) occurred days, weeks, or even months in the past. Thus, we ran another set of models, now focusing only on the Old trials (E2), but with these trials split into two groups based on when the prior exposure occurred (E1). ‘Old-within’ trials corresponded to E2 trials for which E1 occurred within the same session—in other words, memory for the prior exposure was likely to be relatively strong. ‘Old-across’ trials corresponded to E2 trials for which E1 occurred in a prior session (i.e., at least a day in the past)—in other words, memory for the prior exposure was likely to be relatively weak or even absent. Strikingly, for the Old-within trials, there was no effect of global similarity for any of the parietal or VOTC ROIs (*p’s* > 0.14). In contrast, for the Old-across trials there were significant effects of global similarity for each of the ROIs (AnG, *β* = 0.081, *Z* = 3.38, *p* < 0.001; LatIPS, *β* = 0.077, *Z* = 3.23, *p* = 0.001; pIPS, *β* = 0.102, *Z* = 4.24, *p* < 0.001; VOTC, *β* = 0.104, *Z* = 4.28, *p* < 0.001; all survived correction). Moreover, the effect of global similarity was significantly stronger for Old-across trials than Old-within trials in AnG (*Z* = 2.24, *p* = 0.025), LatIPS (*Z* = 2.35, *p* = 0.019) and VOTC (*Z* = 2.20, *p* = 0.027), but not in pIPS (*Z* = 1.53, *p* = 0.126). Thus, when a true memory for past experience with a stimulus was relatively strong (Old-within trials), this substantially reduced the influence of global similarity on memory decisions.

### Tradeoff between global similarity and true memory signals

Our interpretation of the results for the Old-within trials is that the influence of global similarity was reduced by the availability of a true memory for prior experience with the stimulus (E1 memory). To test this prediction more directly, we constructed another set of models—again using the Old-within and Old-across groupings and the same exclusion criteria as in the preceding model—but we now replaced global similarity with a measure of same-stimulus similarity. That is, we simply computed the E1-E2 pattern similarity and used this as a predictor of memory decisions (at E2). Note: for this model, we did not subtract ‘forward’ pattern similarity (E2-E3 similarity) from ‘backward’ pattern similarity (E1-E2 similarity) because the spacing between events was variable. In other words, it was not possible to create symmetrical measures.

The influence of same-stimulus similarity on memory decisions was significant across each of the lateral parietal and VOTC ROIs for the Old-within (AnG, *β* = 0.185, *Z* = 2.49, *p* = 0.013; LatIPS, *β* = 0.386, *Z* = 4.82, *p* < 0.001; pIPS, *β* = 0.490, *Z* = 6.56, *p* < 0.001; VOTC, *β* = 0.506, *Z* = 7.08, *p* < 0.001; all survived correction) and Old-across trials (AnG, *β* = 0.151, *Z* = 5.74, *p* < 0.001; LatIPS, *β* = 0.119, *Z* = 4.38, *p* < 0.001; pIPS, *β* = 0.148, *Z* = 5.01, *p* < 0.001; VOTC, *β* = 0.112, *Z* = 4.03, *p* < 0.001; all survived correction). Specifically, stronger E1-E2 pattern similarity was associated with a higher probability of endorsing an E2 stimulus as ‘old.’ Critically, however, this effect was significantly stronger for Old-within trials than Old-across trials in LatIPS (*Z* = 3.18, *p* = 0.001), pIPS (*Z* = 4.34, *p* < 0.001), and VOTC (*Z* = 5.19, *p* < 0.001); for ANG, there was no significant difference between the trial types (*Z* = 0.43, *p* = 0.667). Thus, the relative recency of E1 had opposite effects on the influence of global similarity vs. same-stimulus similarity: when E1 appeared in the same session as E2, the influence of same-stimulus similarity was relatively greater and the influence of global similarity was relatively lower; in contrast, when E1 appeared in a different session as E2 (further in the past), the influence of same-stimulus similarity was relatively lower and the influence of global similarity was relatively higher. This pattern of data indicates that a strong, ‘true’ memory can override the influence of global similarity; but, in the absence of a strong, ‘true’ memory, global similarity has a powerful influence on memory decisions.

### Activity patterns in parietal cortex reflect high-level / semantic content

While the results above demonstrate that activity patterns in lateral parietal and VOTC ROIs reflected information that was relevant to memory decisions, they do not specify the *nature* of the information in these activity patterns. To address this, we conducted a final set of analyses in which we tested for relationships between activity patterns in each ROI and information content within different layers of a deep convolutional neural network^26–29^. Specifically, we passed the stimuli (natural scene images) through pre-trained VGG-16 model^30^ to obtain the activation pattern for each image at each processing layer. For this analysis, we used the 907 images that were shared across all 8 subjects. Using the VGG-16 activation patterns, we constructed a representational dissimilarity matrix (RDM) by calculating pairwise Pearson correlation between obtained activation patterns for each layer of the model (RDM_VGG-16_). Similarly, for each subject, we then constructed RDMs for each ROI (RDM_Neural_) from fMRI activation patterns. We then performed Spearman correlations between RDM_VGG-16_ and RDM_Neural_ in order to measure the degree to which representational structure in a given brain region resembled the representational structure in a given VGG-16 layer. Statistical significance at the group level was assessed using one-sided Wilcoxon signed-rank tests^31^. We hypothesized that representational structure in lateral parietal and VOTC regions would most closely resemble representational structure in relatively late layers of VGG-16 (which are thought to reflect higher-level, semantic content).

Significant positive correlations between the RDM_VGG-16_ and the RDM_Neural_ were observed across many ROIs. These correlations were observed for relatively late layers in AnG (layers 3-8, all rank sum ≥ 36, *p’s* < 0.004), LatIPS (layer 6-8, all rank sum ≥ 32, *p’s* < 0.027), pIPS (layer 4-8, all rank sum ≥ 34, *p’s* < 0.012), and VOTC (layer 3-8, all rank sum ≥ 35, *p’s* < 0.008). In contrast, for EVC, correlations were strongest in relatively early layers (layer 1-4, all rank sum ≥ 33, *p’s* < 0.020). Significant correlations were also observed in HPC (layer 3-8, all rank sum ≥ 36, *p’s* < 0.004), but not in M1. Of particular relevance, the similarity between VGG-16 and fMRI RDMs increased as a function of VGG-16 model layer in the lateral parietal and VOTC ROIs. Specifically, a Spearman correlation between layers (1 to 8) and RDM similarities revealed a significant positive relationship in AnG (*rho* = 0.64, *p* < 0.001), LatIPS (*rho* = 0.72, *p* < 0.001), pIPS (*rho* = 0.66, *p* < 0.001), and VOTC (*rho* = 0.66, *p* < 0.001). A similar effect was observed in HPC (*rho* = 0.64, *p* < 0.001). In contrast, there was a significant negative relationship between layers (1 to 8) and RDM similarities in EVC (*rho* = −0.84, *p* < 0.001). Together, these results demonstrate a clear distinction between the information tracked by early visual cortex versus lateral parietal and VOTC ROIs. Of central relevance, all of the ROIs in which we observed effects of global similarity on memory decisions (the parietal ROIs and VOTC) were characterized by a preference for higher-level, semantic information. This is consistent with the idea that global similarity effects on memory operated at a relatively high representational level.

**Figure 4.**
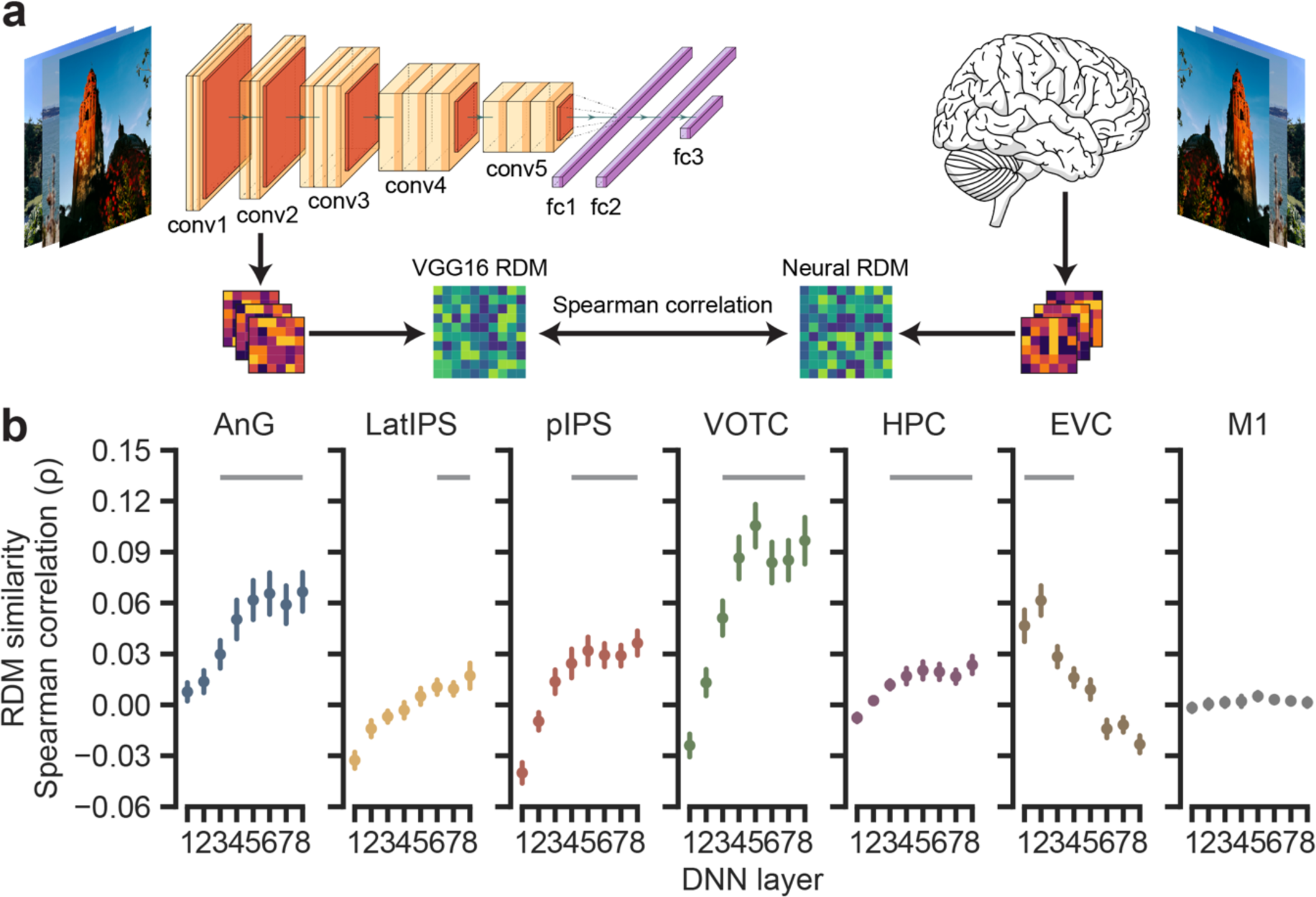
Similarity between fMRI and VGG-16 representations. **a,** A schematic of how representational dissimilarity matrices (RDMs) were calculated. Images that all subjects viewed in the fMRI experiment were passed to the deep neural network (DNN) model (VGG-16). Then the activation patterns of each DNN layer were extracted and pairwise distances (based on Pearson correlations) between images were calculated to form the neural network RDMs. Similarly, fMRI activation patterns were extracted for each of the same images, separately for each ROI and subject, to form the fMRI RDMs. Spearman correlations were then calculated between the neural network RDMs and the fMRI RDMs to quantify the correspondence (similarity) in representations. **b,** Spearman’s rank correlation coefficients between the neural network (VGG-16) RDMs and the fMRI RDMs. The neural network RDMs are separated by DNN layer, which represent different processing stages. Grey bars indicate the layers with significant RDM correlations between the neural network layer and fMRI ROI. Error bar denotes the standard error.

## Discussion

Here, using data from a massive fMRI recognition memory study^15^, and inspired by classic theories in cognitive psychology^7–9^, we show that trial-by-trial recognition memory decisions are predicted by temporally-asymmetric neural measures of global similarity. Specifically, we found that the probability of endorsing a current memory probe as ‘old’ was positively related to the strength of global similarity to past events *relative to* future events. Notably, this relationship was present in regions of lateral parietal cortex that have consistently been implicated in episodic memory^16–18^. Importantly, however, the influence of global similarity on memory decisions depended on the mnemonic history of the probe: global similarity had the strongest influence when the probe was either novel or had initially been encoded at least a day in the past. Finally, using convolutional neural networks, we show that the brain regions in which global similarity predicted memory decisions are regions that preferentially express high-level semantic information, revealing a specific representational level at which similarity-based memory decisions operate.

### Isolating global similarity in time

A unique and critical feature of our analysis approach is that we separately computed global similarity using experiences from the *past* and experiences in the *future*. Our motivation for this approach is that global similarity to future events serves as a baseline that captures time-invariant similarity between a probe and other (normative or typical) experience. Thus, by subtracting future similarity from past similarity, we controlled for generic properties of probes (like typicality) that could lead to higher global similarity values *and* a higher likelihood of ‘old’ decisions. This simple step powerfully isolates the influence that past experience, per se, exerts on current memory decisions. However, our approach begs the question: is this form of baseline correction something the brain actually computes? Our position is that it is sub-optimal for memory decisions to rely on undifferentiated global similarity (this would lead to excessive false recognition). Thus, there is adaptive value in differentiating similarity that arises from recent experience from similarity based on a lifetime of experience. While it is obviously not possible that the brain computes similarity to future events (the specific analysis we employed), the brain could baseline recent experience against any sample of time (e.g., the very distant past) that captures ‘typical experience’. Thus, other variants of our analysis that used the distant past instead of the future would be conceptually equivalent. Here, however, we used future events as a baseline because it captured normative experience but is ‘completely free’ from episodic memory.

In addition to comparing similarity to past vs. future events, we also sampled from different temporal windows in the past. This sampling of temporal windows was a unique feature of our analysis approach that was only enabled by the use of a recognition memory task that spanned many fMRI sessions. By comparing global similarity across different temporal windows, we were able to test a straightforward, but important prediction: that events from the distant past exert relatively less influence on current memory decisions than events from the recent past. To the extent that older memories are weaker^32^, this represents another way to confirm that any observed relationship between global similarity and memory decisions is based on the influence of actual memories, as opposed to more generic properties of a probe. Notably, global matching models were originally developed and applied to explain memory decisions in paradigms where a single list of studied materials (e.g., words) was followed by a single test list (probes)^5,9^. In these paradigms, global matching models ignored the recency of past experience—instead, all items from the study list were given equal weight on memory decisions in the test list. In more recent work, forgetting or decay has been included as a parameter in global matching models^33^ in order to ‘de-weight’ older memories. That said, prior work has not explicitly considered or quantified the influence of past events on current decisions as a function of their temporal recency.

Interestingly, we did not find evidence that past experience influenced current memory decisions in the immediate past condition (<1 minute in the past). We believe this null result should be interpreted with caution because the immediate past condition averaged over fewer trials (by definition, we were sampling a narrower time window) and it involved correlating trials from the same scan run as the probe, raising potential concern about non-independence (autocorrelation) between the probe trial and immediately preceding trials (though, in principle, the backward – forward global similarity measure should control for effects of autocorrelation). That said, a potential cognitive account of this null effect is that events from the immediate past are retained at a higher fidelity in memory and, therefore, it is easier to differentiate these events from a current memory probe. Thus, while the null effect for the immediate past condition represents an interesting observation that could be explored in a more targeted manner in future studies, this was not an *a priori* prediction and it is not relevant to our core conclusions.

### Implications for global matching models

While our analytic approach was directly inspired by classic global matching models, it is important to emphasize that there are many variants of, and parameters within, these models. Here, our goal was not to systematically compare these variants and parameters to arrive at an optimal model; rather, we used a form of these models as a tool for identifying neural measures that reflected the influence of past experience on current memory decisions. However, one important test we did include was to compare global similarity (which averages across many trials) to ‘maximum similarity’—that is, the highest similarity between a probe and an event from the past. Critically, global similarity markedly outperformed maximum similarity in predicting trial-by-trial memory decisions, confirming that there is an advantage to considering all experiences from a given temporal window.

Our analyses also reveal an important and striking caveat to the relationship between global similarity and memory decisions: this influence is substantially reduced when a probe’s prior experience (E1) is readily available in memory. Specifically, by focusing on probes that were veridically old (E2 trials), we were able to compare the influence of global similarity on memory decisions as a function of whether E1 occurred within the same experimental session or in an experimental session days to months ago. Whereas global similarity robustly predicted memory decisions when E1 had been studied at least a day in the past (across sessions), there was no influence of global similarity when E1 had been studied in the same session (and E1 memory was presumably much stronger). This dissociation was paralleled by a dramatic and opposite shift in the influence of same-stimulus similarity (E1-E2 similarity) on memory decisions. Namely, when E1 occurred in the same session as E2, the relationship between E1-E2 similarity and memory decisions was much *stronger* compared to when E1 had occurred in a prior session. Taken together, this pattern of data reveals a clear tradeoff between global similarity and same-stimulus similarity. When memory for a prior occurrence of an event (E1) is weak, then global similarity drives memory decisions, but when memory for a prior occurrence of an event is strong, same-stimulus similarity dominates.

### Brain regions in which global similarity predicted memory decisions

Our a priori interest in lateral parietal cortex (LPC) was motivated by substantial evidence implicating LPC in recognition memory decisions^18,34–37^. However, understanding the role of LPC in memory has been a subject of much debate. One key line of evidence that has helped constrain theories of LPC contributions to memory is that LPC actively represents the *contents* of memories^19–21,38^. Our findings are consistent with this literature, but also constitute an important advance in that, here, we explicitly link LPC content representations—from specific temporal windows in the past—to trial-by-trial recognition memory decisions^10–12^. The fact that memory decisions were predicted by LPC content representations across a timescale of minutes is reminiscent of evidence—outside the domain of memory—which has described LPC as having a wide ‘temporal receptive window.’ Specifically, LPC—and angular gyrus, in particular—has been shown to integrate information across relatively long timescales—on the order of minutes. Thus, an account of the current findings that bridges across these literatures is that LPC is able to integrate content across relatively long timescales and these integrated content representations could potentially support everything from following a story^39,40^ to recognition memory decisions.

The idea of temporal integration does raise an interesting question: does global similarity reflect a memory search process initiated by the probe, or does the brain compute a running average of experience (i.e., an integrated representation) that is automatically compared to the probe—or even serves as a prediction of upcoming experience? These ideas, which have a precedent in the decision-making literature^41^, could be tested by determining whether the *relationship* between global similarity and memory decisions is influenced by top-down (memory search) goals. For example, if the relationship between global similarity and memory decisions is influenced by instructions to search within specific temporal windows (e.g., “Did you see this stimulus *yesterday*?” vs. “Did you see this stimulus *today*?”), this would strongly favor a search account. In contrast, if the relationship between global similarity and memory decisions is not influenced by such goals (even if subjects can use these goals to constrain responses), then this would strongly favor a running average account.

To more definitively and precisely establish content representations within the LPC regions that showed global similarity effects, we used VGG-16 (a deep convolutional neural network) to measure content effects across different network layers. We found that the regions that demonstrated relationships between global similarity and memory decisions (LPC and ventral temporal cortex) were characterized by markedly stronger representations of information at late VGG-16 layers. These late layers are thought to represent high-level or semantic information, as opposed to early layers which capture lower-level visual properties. Indeed, the pattern of data in LPC and ventral temporal cortex contrasted sharply with early visual cortex, where early layers were preferentially represented. Thus, our findings not only implicate LPC in reflecting global similarity, but indicate that the specific representational level of similarity in these regions—and the representations that putatively drive memory decisions—is related to high-level semantic information^42,43^.

Notably, we did not observe any relationship between global similarity in the hippocampus and recognition memory decisions. While there is a robust literature implicating the hippocampus in episodic memory, our analysis approach focused on global similarity averaged across many stimuli—a measure that is potentially misaligned with the computations the hippocampus supports. Indeed, prior evidence specifically highlights a dissociation between global similarity measures in neocortical areas versus more stimulus-specific representations in the hippocampus^44^. Interestingly, some variants of global matching models have applied nonlinear transformations (e.g., cubic or exponential) to global similarity values in order to more strongly weight the influence of highly similar matches^4,6^. While beyond the scope of the current manuscript, it is possible that with the right parameters, global matching models may better ‘fit’ the computations that the hippocampus supports.

## Conclusions

Using an innovative analysis approach and a highly unique dataset, we show that trial-by-trial memory decisions are predicted by temporally-asymmetric neural measures of global similarity. These measures of global similarity were robustly expressed in regions of lateral parietal cortex that tracked high-level semantic content. Together, these results provide a new framework for measuring and conceptualizing the neural computations that support recognition memory.

## Methods

All analyses described here were based on a previously-published and extensively characterized dataset: the Natural Scenes Dataset (NSD)^15^. Relevant details, including unique statistical analyses, are described below.

### Subjects

Eight subjects (six female, mean age = 26.5 years, range = 19 – 32 years) participated in the experiment. All subjects had normal or corrected-to-normal vision. Written consent was obtained from all subjects. The study was approved by the University of Minnesota Institutional Review Board.

### Stimuli and experimental procedure

All stimuli used in the experiment were selected from Microsoft’s COCO image database (Lin et al., 2014). A set of 73,000 colored images were selected from 80 categories, out of the 90 original COCO categories. Images were cropped into square (425×425 pixels). A screening procedure was implemented to remove duplicate, extremely similar, or potentially offensive images. In the experiment, subjects performed a long-term continuous recognition task. It was intended that each subject would view 10,000 unique images, each repeated 3 times, distributed over 40 fMRI sessions. Out of the 10,000 images, 1,000 of them were shared across all subjects and the remaining 9,000 were unique to each subject. During each trial, an image was presented on screen for 3 seconds, followed by a 1 second blank screen. Subjects were instructed to press one of two buttons to indicate whether the image had been presented at any prior point in the experiment (including in prior sessions; ‘old’) or was novel (‘new’). Thus, for every trial in the experiment, the current stimulus served as a ‘probe’ that was to-be-compared against all previously-studied stimuli. Subjects were additionally instructed to fixate a central dot throughout the entire task.

Within each fMRI session, there were 12 runs of the continuous recognition task that displayed a total of 750 natural scene images. Each run lasted 300s and contained 75 trials. The first 3 and the last 4 trials were blank trials. For odd-numbered runs, the remaining 68 trials consisted of 63 stimulus trials and 5 randomly-distributed blank trials. For even-numbered runs, the remaining 68 trials consisted of 62 stimulus trials, 5 randomly-distributed blank trials, and one ‘fixed’ blank trial (trial #63). While each subject studied a (mostly) unique set of images, the distribution of image exposures (E1, E2, E3) across the 40 sessions had an identical structure for each subject in order to minimize differences in recognition memory performance. E1, E2, and E3 trials were distributed across all 40 sessions but the proportion of these trials changed across sessions: from E1 = 77.7%, E2 = 18.1%, and E3 = 4.2% in session 1 to E1 = 4.0%, E2 = 19.7%, and E3 = 76.3% in session 40. Within each session, for each E1 trial, there was a 43.8% (±18.4%) chance on average that the image would repeat within the session (E2); otherwise, corresponding E2 trials were uniformly distributed across the remaining sessions. Note: not all subjects finished all 40 fMRI sessions (range was 30 – 40 sessions). To minimize across-subject differences, here we only analyzed data from the first 30 sessions for each subject.

### MRI acquisition

MRI data were collected at the Center for Magnetic Resonance Research at the University of Minnesota. Functional data and fieldmaps were collected using a 7T Siemens Magnetom passively shielded scanner with a 32 channel head coil. A gradient-echo EPI sequence at 1.8mm isotropic resolution with whole brain coverage was used to acquire functional data (84 axial slices, slice thickness = 1.8mm, slice gap = 0mm, field-of-view = 216mm × 216mm, phase-encode direction A-P, matrix size = 120 × 120, TR = 1600ms, TE = 22.0ms, flip angle = 62°, echo spacing = 0.66ms, partial Fourier = 7/8, in-plane acceleration factor (iPAT) = 2, multiband acceleration factor = 3). Several dual-echo EPI fieldmaps were acquired periodically over each scan session (2.2mm × 2.2mm × 3.6mm resolution, TR = 510ms, TE1 = 8.16ms, TE2 = 9.18ms, flip angle = 40°, partial Fourier = 6/8). Anatomical images were collected using a 3T Siemens Prisma scanner with a standard 32 channel head coil. Several (6 – 10) whole brain T_1_-weighted scans were acquired for each subject across the experiment using an MPRAGE sequence (0.8mm isotropic resolution, TR = 2400ms, TE = 2.22ms, TI = 1000ms, flip angle = 8°, in-plane acceleration factor (iPAT = 2). In addition, several T_2_-weighted scans were obtained using a SPACE sequence (0.8mm isotropic resolution, TR = 3200ms, TE = 563ms, in-plane acceleration factor (iPAT) = 2 to facilitate medial temporal lobe subregion identification.

### MRI data processing

All the pre-processed data were taken directly from the Natural Scenes Dataset; pre-processing steps are described in detail in the data paper^15^. In brief, T_1_-weighted and T_2_-weighted images were corrected for gradient nonlinearities using the Siemens gradient coefficient file from the scanner. All T_1_ and T_2_ images for a given subject were co-registered to the 1st T_1_ volume. The final version of T_1_ and T_2_ images were resampled from the co-registered data using cubic interpolation to 0.5mm isotropic resolution. Finally, the multiple images within each modality were averaged to improve signal to noise ratio. The averaged T_1_ image was processed by FreeSurfer 6.0.0 with *-hires* option enabled. Manual edits were performed to improve the accuracy of surface reconstruction. Utilizing surfaces generated by FreeSurfer, several additional cortical surfaces between the pial and white matter were generated at 25%, 50%, and 75% cortical depth. These surfaces were used to map the volume data to surface space. For fMRI data, all pre-processing was performed in the subjects’ native space. Images were first corrected for slice-timing and upsampled to 1s. Then gradient nonlinearities, spatial distortion and motion correction were performed. fMRI images from later NSD sessions were co-registered to the mean fMRI volume of the first NSD session. All the spatial transformations were concatenated to allow a single step cubic interpolation. In this step, data was upsampled to 1mm isotropic resolution.

To model the neural responses of each trial, a GLM was fitted for each NSD session using the package GLMsingle^46^. Optimal HRFs were chosen for each voxel from a library of HRFs to better compensate for differences in hemodynamic responses. Each trial was modeled separately in the model using the optimal HRF. The detailed procedure of this method is described in Allen et al.^15^ and the results denoted as ‘b2’ version in the paper. Models were fitted on the pre-processed fMRI data in 1mm functional space. The estimated single-trial betas were further resampled to each of the three cortical surface depths and averaged together using cubic interpolation. The result was then transformed to fsaverage space using nearest neighbor interpolation. This version of betas was used in analyses of cortical regions.

### Regions of interest

ROIs were defined in fsaverage space (cortical regions) and the subjects’ native 1mm functional space (hippocampus). For all cortical ROIs, we used the multi-modal parcellation (MMP1)^47^. Based on previous related studies^10–12^, we focused our main analyses on lateral parietal cortex and subdivided it into the angular gyrus (AnG), lateral intraparietal sulcus (LatIPS) and posterior intraparietal sulcus (pIPS) regions. We combined MMP1 label PGs and PGi regions to create AnG. For LatIPS and pIPS, we combined regions to match the definition used in Favila et al. ^19^ as closely as possible. Namely, the LatIPS consisted of MMP1 label IP1, IP2 and LIPd. The pIPS consisted of MMP1 labels IP0, IPS1, MIP, VIP and LIPv. In addition, we also included ventral occipitotemporal cortex (VOTC) and hippocampus (HPC), given the involvement of these two regions in memory process. The VOTC consisted of MMP1 labels FFC, VVC, PHA1, PHA2, PHA3, PIT, V8, VMV1, VMV2 and VMV3. The HPC used a manually traced segmentation in subject’s native space that combined subregions CA1, CA2, CA3, dentate gyrus and hippocampus tail. The early visual cortex (EVC, MMP1 label: V1) and premotor cortex (M1, MMP1 label: Area4) were also included as control ROIs. All ROIs were combined across the left and right hemispheres.

### Neural measures of global similarity

To compute neural measures of global similarity, we compared the fMRI pattern evoked by a ‘current stimulus’ (probe) to activity patterns evoked by past and future trials. However, because of the continuous recognition design, a given trial potentially served in all three roles (probe, past, future) as the analyses were iteratively performed (trial-by-trial). Thus, probes were not separate trials, but instead a designation of the trial’s role in a particular iteration of an analysis.

For each probe, we constructed three temporal windows representing past experience: Immediate, Recent, and Distant. The *immediate temporal window* included the past 15 trials within the same scan run (mean temporal distance to current trial = 35.0 seconds, range: 4.0 seconds to 68.0 seconds), the *recent temporal window* included trials from the past 3 scan runs (mean = 3.9 minutes, range: 2.8 minutes to 37.1 minutes), and the *distant temporal window* included trials from the prior fMRI session (mean = 7.3 days, range: 1.0 days to 28.0 days). Mirror-reversed, but otherwise identical temporal windows were also constructed for future experience.

Importantly, a given trial was only included as a probe if it allowed for *all* of the temporal windows to be constructed. Thus, probes never ‘occurred’ in the first or last session (session 1 or session 30), in the first or last three runs within a session, or in the first 15 or last 15 trials in a run. For example, trial #14 in a given scan run was never included as a probe in any analysis because the immediate temporal window could not be constructed (there were not 15 preceding trials). Thus, even though each temporal window imposed different constraints, we only included a trial as a probe if it met the constraints for each of the temporal windows. However, even if a trial was excluded as a probe, it could serve *within* a temporal window. For example, trial #14 would be part of the recent past temporal window for trial #16, assuming it did not occur in the first or last session or the first or last three runs within a session. Trials were also excluded as probes if no behavioral response was made on that trial.

For each probe, we computed the Pearson correlation between the activity pattern evoked on that trial and each trial within each temporal window from the past and future. These correlation values were then Fisher’s *Z*-transformed and averaged within each temporal window, separately for past and future. Finally, for each probe and each temporal window, we subtracted the ‘forward’ similarity (similarity to future events) from the ‘backward’ similarity (similarity to past events). This yielded, for each trial and each temporal window (immediate, recent, distant), a difference score which served as the measure of global similarity. Thus, values of 0 represented no difference in mean similarity between past and future, values greater than 0 represented relatively higher similarity to the past, and values below 0 represented relatively higher similarity to the future. This entire process was separately performed for each ROI. Note: unless noted otherwise, a probe’s temporal window could potentially contain a repetition of the same stimulus as the probe. For example, if a probe was an E2 trial, the corresponding E1 trial might fall within one of the three temporal windows in the past. For the control analysis using maximal similarity, the criteria for selecting the probes were identical. However, instead of averaging similarity values across all trials in a temporal window, we identified the single trial, within each temporal window, with the highest similarity value. We then subtracted the highest value from each future temporal window from the highest value from the corresponding past temporal window. For analyses based on *same-stimulus similarity*, although temporal windows were not relevant, for consistency we retained the same criteria for selecting probes as in the global similarity and maximal similarity analyses, with the exception that only E2 trials (the 2^nd^ presentation of a stimulus) served as probes. Same-stimulus similarity was calculated as the Fisher’s *Z*-transformed Pearson correlation between the fMRI activation pattern evoked by the probe (E2) and the corresponding E1 trial. In this analysis, we did not subtract ‘forward’ pattern similarity (E2-E3 similarity) from ‘backward’ pattern similarity (E1-E2 similarity) because the distance between E1 and E2 was not matched with the distance between E2 and E3; moreover, this was not relevant, conceptually, for this analysis.

### Representational similarity matrices from fMRI and neural networks

Separate representational dissimilarity matrices (RDMs) were constructed based on fMRI data and a deep convolutional neural network. These RDMs were restricted to images that were shared across all eight subjects and, to match the main analyses, only to images that were presented during the first 30 NSD sessions. This resulted in a total of 907 images that were used for the RDMs. For the fMRI-based RDMs, activity patterns for each trial within an ROI were extracted, then averaged across exposures for each image, resulting in a single, averaged pattern per image and ROI. Then, pairwise Pearson correlations (Fisher’s *Z* transformed) were calculated for each pair of images, yielding an RDM for each ROI. For the neural network RDM, we utilized a pre-trained version of VGG-16^30^ included with the *torchvision* package (https://github.com/pytorch/vision). For each image, the same preprocessing steps (resizing, intensity normalization) were applied as used for the images in VGG16 training. The preprocessed images were passed to the model and, for each image, the unit activation at each processing layer served as the image ‘representation’. Specifically, eight layers of activation were used in the RDM similarity analyses, which corresponded to 5 pooling layers (2^nd^, 4^th^, 7^th^, 10^th^, 13^th^) and three fully connected layers. These layers were labeled as layer 1 to 8 as they progressed in the processing hierarchy of VGG-16. Similar to the fMRI-based RDM, pairwise Pearson correlations (Fisher’s *Z* transformed) were calculated for each pair of images to form RDMs for each layer. Spearman correlations were calculated between fMRI and VGG-16 based RDMs to quantify the similarity between the fMRI and neural network representations.

### Statistical analyses

#### Logistic mixed effects models

Logistic mixed effects models were used to model the relationship between global similarity and subjects’ memory responses (old vs. new). The main model included global similarity from 3 temporal windows (immediate, recent, distant; each window representing backward – forward similarity); an image’s exposure/repetition number (1^st^, 2^nd^, 3^rd^); and the interaction between these variables as fixed effects. In addition, to account for subjects’ potential response biases and session effects, the proportion of a subject’s old responses within each temporal window and the NSD session number were added in the model as fixed effect confound regressors. Subject ID was included as a random effect with random intercept only. The model formula was:

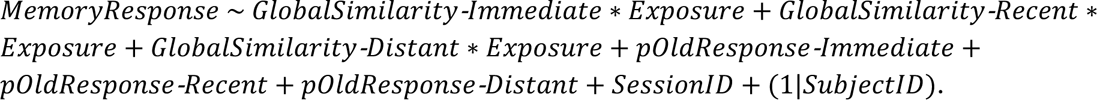

For the models examining maximum similarity, the formula was the same as the main model, except global similarity was replaced with the maximum pattern similarity (backward – forward) within each temporal window.

To examine the effect of global similarity on memory decisions for Old vs. Novel probes, we constructed a new set of models that focused only on the recent temporal window and only included probes corresponding to E1 (1^st^ exposure; Novel) or E2 (2^nd^ exposure; Old). All E3 trials were excluded so that the number of Novel and Old trials was relatively balanced. Importantly, for this set of analyses we also excluded any E2 trials for which the corresponding E1 trial fell within the recent temporal window. The model formula was:

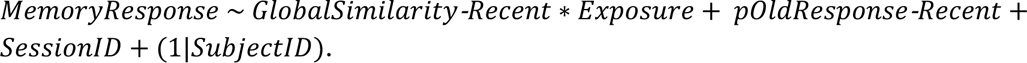

To test whether the temporal lag between E1 and E2 influenced the relationship between global similarity and memory decisions, we constructed a separate set of models that only included probes corresponding to E2 trials, but with these trials split into two conditions based on when the prior exposure (E1) occurred. ‘Old-within’ corresponded to trials for which E1 occurred within the same experimental session as E2. ‘Old-across’ corresponded to trials for which E1 occurred in a prior experimental session. The model formula was:

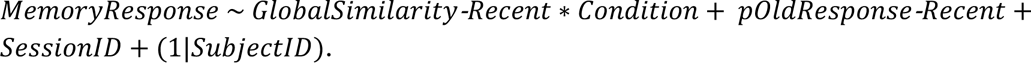

Finally, for the models that tested for relationships between same-stimulus similarity and memory decisions, the models were identical to the preceding set of models except that global similarity was replaced with same-stimulus similarity.

All models were fit using the package *lme4* (https://cran.r-project.org/web/packages/lme4/index.html) in *R*. Likelihood ratio tests were used to determine the significance of fixed effects. For post-hoc tests of fixed effects and interactions, Wald test with asymptotic distribution was used with package *emmeans* (https://cran.r-project.org/web/packages/emmeans/index.html) in *R*.

#### RDM similarity

Spearman’s rank correlation was used to quantify the similarity between the fMRI and neural network (VGG-16) RDMs. The correlation coefficients were calculated within each subject. One-sided Wilcoxon signed-rank tests were used to determine the significance at the group level^31^. For comparison of different VGG-16 layers, two-sided Wilcoxon signed-rank tests were used.

## Acknowledgements

This work was supported by NIH-NINDS R01NS089729 and NIH-NINDS R01NS107727 to B.A.K and by NIH-NEI R01EY023384 to T.N. Collection of the Natural Scenes Dataset was supported by NSF IIS-1822683 (K.K.) and NSF IIS-1822929 (T.N.).

## Reference

1. Mandler, G. Recognizing: The judgment of previous occurrence. Psychol. Rev. 87, 252–271 (1980).

2. Suzuki, W. A. & Eichenbaum, H. The Neurophysiology of Memory. Ann. N. Y. Acad. Sci. 911, 175–191 (2006).

3. Yonelinas, A. P. The Nature of Recollection and Familiarity: A Review of 30 Years of Research. J. Mem. Lang. 46, 441–517 (2002).

4. Hintzman, D. L. Judgments of frequency and recognition memory in a multiple-trace memory model. Psychol. Rev. 95, 528–551 (1988).

5. Murdock, B. B. A theory for the storage and retrieval of item and associative information. Psychol. Rev. 89, 609–626 (1982).

6. Nosofsky, R. M. Attention, similarity, and the identification–categorization relationship. J. Exp. Psychol. Gen. 115, 39–57 (1986).

7. Arndt, J. & Hirshman, E. True and False Recognition in MINERVA2: Explanations from a Global Matching Perspective. J. Mem. Lang. 39, 371–391 (1998).

8. Clark, S. E. & Gronlund, S. D. Global matching models of recognition memory: How the models match the data. Psychon. Bull. Rev. 3, 37–60 (1996).

9. Gillund, G. & Shiffrin, R. M. A retrieval model for both recognition and recall. Psychol. Rev. 91, 1–67 (1984).

10. Wing, E. A. et al. Cortical Overlap and Cortical-Hippocampal Interactions Predict Subsequent True and False Memory. J. Neurosci. 40, 1920–1930 (2020).

11. Ye, Z. et al. Neural Global Pattern Similarity Underlies True and False Memories. J. Neurosci. 36, 6792–6802 (2016).

12. Zhu, B. et al. Multiple interactive memory representations underlie the induction of false memory. Proc. Natl. Acad. Sci. 116, 3466–3475 (2019).

13. Bartlett, J. C., Hurry, S. & Thorley, W. Typicality and familiarity of faces. Mem. Cognit. 12, 219–228 (1984).

14. Vokey, J. R. & Read, J. D. Familiarity, memorability, and the effect of typicality on the recognition of faces. Mem. Cognit. 20, 291–302 (1992).

15. Allen, E. J. et al. A massive 7T fMRI dataset to bridge cognitive neuroscience and artificial intelligence. Nat. Neurosci. 25, 116–126 (2022).

16. Hutchinson, J. B. et al. Functional Heterogeneity in Posterior Parietal Cortex Across Attention and Episodic Memory Retrieval. Cereb. Cortex 24, 49–66 (2014).

17. Vilberg, K. L. & Rugg, M. D. Memory retrieval and the parietal cortex: A review of evidence from a dual-process perspective. Neuropsychologia 46, 1787–1799 (2008).

18. Wagner, A. D., Shannon, B. J., Kahn, I. & Buckner, R. L. Parietal lobe contributions to episodic memory retrieval. Trends Cogn. Sci. 9, 445–453 (2005).

19. Favila, S. E., Samide, R., Sweigart, S. C. & Kuhl, B. A. Parietal Representations of Stimulus Features Are Amplified during Memory Retrieval and Flexibly Aligned with Top-Down Goals. J. Neurosci. 38, 7809–7821 (2018).

20. Kuhl, B. A. & Chun, M. M. Successful Remembering Elicits Event-Specific Activity Patterns in Lateral Parietal Cortex. J. Neurosci. 34, 8051–8060 (2014).

21. Xiao, X. et al. Transformed Neural Pattern Reinstatement during Episodic Memory Retrieval. J. Neurosci. 37, 2986–2998 (2017).

22. Grill-Spector, K. & Weiner, K. S. The functional architecture of the ventral temporal cortex and its role in categorization. Nat. Rev. Neurosci. 15, 536–548 (2014).

23. Burgess, N., Maguire, E. A. & O’Keefe, J. The Human Hippocampus and Spatial and Episodic Memory. Neuron 35, 625–641 (2002).

24. Tulving, E. & Markowitsch, H. J. Episodic and declarative memory: Role of the hippocampus. Hippocampus 8, 198–204 (1998).

25. Osth, A. F. & Dennis, S. Global matching models of recognition memory. Preprint at 10.31234/osf.io/mja6c (2020).

26. Guclu, U. & van Gerven, M. A. J. Deep Neural Networks Reveal a Gradient in the Complexity of Neural Representations across the Ventral Stream. J. Neurosci. 35, 10005–10014 (2015).

27. Khaligh-Razavi, S.-M. & Kriegeskorte, N. Deep Supervised, but Not Unsupervised, Models May Explain IT Cortical Representation. PLoS Comput. Biol. 10, e1003915 (2014).

28. Mehrer, J., Spoerer, C. J., Jones, E. C., Kriegeskorte, N. & Kietzmann, T. C. An ecologically motivated image dataset for deep learning yields better models of human vision. Proc. Natl. Acad. Sci. 118, e2011417118 (2021).

29. Schrimpf, M. et al. Brain-Score: Which Artificial Neural Network for Object Recognition is most Brain-Like? Preprint at 10.1101/407007 (2018).

30. Simonyan, K. & Zisserman, A. Very Deep Convolutional Networks for Large-Scale Image Recognition. Preprint at http://arxiv.org/abs/1409.1556 (2015).

31. Nili, H. et al. A Toolbox for Representational Similarity Analysis. PLoS Comput. Biol. 10, e1003553 (2014).

32. Wixted, J. T. & Ebbesen, E. B. On the Form of Forgetting. Psychol. Sci. 2, 409–415 (1991).

33. Kahana, M. J., Zhou, F., Geller, A. S. & Sekuler, R. Lure similarity affects visual episodic recognition: Detailed tests of a noisy exemplar model. Mem. Cognit. 35, 1222–1232 (2007).

34. Gonzalez, A., et al. Electrocorticography reveals the temporal dynamics of posterior parietal cortical activity during recognition memory decisions. Proc. Natl. Acad. Sci. 112, 11066–11071 (2015).

35. Hutchinson, J. B., Uncapher, M. R. & Wagner, A. D. Increased functional connectivity between dorsal posterior parietal and ventral occipitotemporal cortex during uncertain memory decisions. Neurobiol. Learn. Mem. 117, 71–83 (2015).

36. Rutishauser, U., Aflalo, T., Rosario, E. R., Pouratian, N. & Andersen, R. A. Single-Neuron Representation of Memory Strength and Recognition Confidence in Left Human Posterior Parietal Cortex. Neuron 97, 209–220.e3 (2018).

37. Sestieri, C., Shulman, G. L. & Corbetta, M. The contribution of the human posterior parietal cortex to episodic memory. Nat. Rev. Neurosci. 18, 183–192 (2017).

38. Bonnici, H. M., Richter, F. R., Yazar, Y. & Simons, J. S. Multimodal Feature Integration in the Angular Gyrus during Episodic and Semantic Retrieval. J. Neurosci. 36, 5462–5471 (2016).

39. Baldassano, C. et al. Discovering Event Structure in Continuous Narrative Perception and Memory. Neuron 95, 709–721.e5 (2017).

40. Cabeza, R., Ciaramelli, E. & Moscovitch, M. Cognitive contributions of the ventral parietal cortex: an integrative theoretical account. Trends Cogn. Sci. 16, 338–352 (2012).

41. Bornstein, A. M., Khaw, M. W., Shohamy, D. & Daw, N. D. Reminders of past choices bias decisions for reward in humans. Nat. Commun. 8, 15958 (2017).

42. Humphreys, G. F., Lambon Ralph, M. A. & Simons, J. S. A Unifying Account of Angular Gyrus Contributions to Episodic and Semantic Cognition. Trends Neurosci. 44, 452–463 (2021).

43. Lee, H., Keene, P. A., Sweigart, S. C., Hutchinson, J. B. & Kuhl, B. A. Adding meaning to memories: How parietal cortex combines semantic content with episodic experience. J. Neurosci. JN-RM-1919-22 (2023) doi:10.1523/JNEUROSCI.1919-22.2023.

44. LaRocque, K. F. et al. Global Similarity and Pattern Separation in the Human Medial Temporal Lobe Predict Subsequent Memory. J. Neurosci. 33, 5466–5474 (2013).

45. Lin, T.-Y. et al. Microsoft COCO: Common Objects in Context. in Computer Vision – ECCV 2014 (eds. Fleet, D., Pajdla, T., Schiele, B. & Tuytelaars, T.) vol. 8693 740–755 (Springer International Publishing, Cham, 2014).

46. Prince, J. S., et al. GLMsingle: A Toolbox for Improving Single-Trial fMRI Response Estimates. http://biorxiv.org/lookup/doi/10.1101/2022.01.31.47843 (2022) doi:10.1101/2022.01.31.478431.

47. Glasser, M. F. et al. A multi-modal parcellation of human cerebral cortex. Nature 536, 171–178 (2016).

